# Expanding TheCellMap.org to visualize a genome-scale genetic interaction network for a human cell line

**DOI:** 10.64898/2026.03.16.712129

**Authors:** Ira Horecka, Matej Ušaj, Myra P.D. Masinas, Henry N. Ward, Xiang Zhang, Arshia Z. Hassan, Maximilian Billmann, Hannes L. Röst, Chad L. Myers, Michael Costanzo, Brenda Andrews, Charles Boone

## Abstract

Genetic interaction networks map functional connections between genes and their corresponding pathways and complexes. We previously developed TheCellMap.org as a central repository for storing and analyzing quantitative genetic interaction data produced by genome-scale Synthetic Genetic Array (SGA) analysis in the budding yeast, *Saccharomyces cerevisiae*. We have expanded TheCellMap.org to include ~89,000 quantitative genetic interactions identified from genome-scale CRISPR-based analysis of ~4 million human gene pairs in the haploid cell line, HAP1. TheCellMap.org enables users to readily access, visualize and explore human HAP1 genetic interactions, as well as to extract and reorganize sub-networks, applying data-driven network layouts in an intuitive and interactive manner.

## Introduction

Genetic interactions reveal functional relationships between genes and identify genetic modifiers that shape the genotype to phenotype relationship (Costanzo *et al*. 2019). Negative interactions, such as synthetic lethal or sick interactions, arise when a double mutant exhibits a fitness defect more severe than expected based on the combined effects of the single mutants. These interactions typically connect functionally related genes that converge on the same essential biological process (Costanzo *et al*. 2010; Costanzo *et al*. 2016) and are of particular interest because they can be leveraged for developing synthetic lethal cancer therapies (Hartwell *et al*. 1997; Setton *et al*. 2021; Previtali *et al*. 2024). In contrast, positive genetic interactions occur when a double mutant displays a less severe phenotype than predicted by the single mutants (Costanzo *et al*. 2019). Genetic suppression is an important, though relatively rare, class of positive interaction in which the double mutant exhibits higher fitness than the sickest single mutant (van Leeuwen *et al*. 2016; Costanzo *et al*. 2019; van Leeuwen *et al*. 2020; Unlu *et al*. 2023). If suppression of a disease gene occurs through a loss-of-function allele, it highlights the potential for the development of an inhibitory small molecule as a possible therapeutic strategy (Chen *et al*. 2016; Masud *et al*. 2025).

Despite their mechanistic differences, both negative and positive genetic interactions are highly organized such that functionally related genes often display similar patterns of genetic interactions (Costanzo *et al*. 2019). The genetic interaction profile of a given gene is made up of its collection of negative and positive genetic interactions. These profiles provide a quantitative readout of gene function, such that genes with highly similar profiles often function in the same biological process, pathway or complex (Costanzo *et al*. 2019). Previous large-scale analyses in the budding yeast, *Saccharomyces cerevisiae*, mapped a global network of ~1,000,000 negative and positive interactions among its ~5,000 nonessential and ~1,000 essential genes (Costanzo *et al*. 2010; Costanzo *et al*. 2016) and this dataset is publicly accessible through TheCellMap.org (Usaj *et al*. 2017).

In a recent study, we constructed a large-scale genetic interaction network for the haploid human cell line, HAP1. We first constructed query mutant cell lines that carry a loss-of-function allele of a specific gene of interest and then tested them for genetic interactions through genome-wide CRISPR screens that target ~17,000 human genes. In total, we examined ~4 million gene pairs and scoring ~89,000 genetic interactions, including ~47,000 negative and ~42,000 positive genetic interactions (Billmann et al., 2026; Cell, *in press*)(Billmann *et al*. 2025b). This work demonstrated that the core organizational properties of genetic networks are broadly conserved from yeast to human cells. Like the global yeast network (Costanzo *et al*. 2016), the HAP1 network of genetic interaction profile similarities organizes genes into a hierarchical model of cellular function. At the most refined resolution, genes cluster into modules corresponding to protein complexes and pathways. At an intermediate level, these modules are merged to define discrete biological processes. At a more general level, clusters representing distinct biological processes group into larger modules corresponding to major cellular compartments (Billmann *et al*. 2025b). The HAP1 network provides a resource for human gene function prediction and supports the systematic functional annotation of independent gene sets. For example, we linked ~100 genes associated with relatively few citations and/or GO annotations to specific bioprocesses based on their location on the HAP1 genetic interaction profile similarity network (Billmann *et al*. 2025b). Here, we describe the expansion of TheCellMap.org, a web-accessible database and visualization platform, designed to facilitate access and exploration of the HAP1 genetic interaction network.

## Materials and Methods

### HAP1 genetic interaction analysis

Genome-wide CRISPR-Cas9 genetic interaction screens, using the pooled lentiviral TKOv3 guide RNA (gRNA) library, were conducted as described elsewhere (Billmann *et al*. 2025a; Billmann *et al*. 2025b).

### Database development

TheCellMap.org is an interactive web application that enables efficient querying of large network datasets. The backend is implemented in Python (https://python.org) using the Django (https://www.djangoproject.com) web framework, connected to a dual-database system comprising a PostgreSQL (https://www.postgresql.org) object-relational database for structured data storage and querying, combined with rasdaman for multi-dimensional array data management (https://doi.org/10.5281/zenodo.1163021). The front-end is built with React (https://reactjs.org), a JavaScript library for creating user interfaces, utilizing JSX to define dynamic components, with Cascading Style Sheets (CSS) (https://www.w3.org/TR/css) styling for an enhanced user experience. Sigma.js (http://sigmajs.org) is integrated into the front-end to render and interactively visualize the genetic interaction network. A nginx (https://nginx.org) web server serves the intermediary between client and server sides, handling static file delivery and routing Hypertext Transfer Protocol (HTTP) (https://httpwg.org) requests through a Web Server Gateway Interface (uWSGI) (https://uwsgi-docs.readthedocs.io). All computationally intensive requests are processed server-side in Python using numpy for rapid numerical analysis of queried interaction data, thus reducing client-side processing overhead and optimizing browser performance (Harris *et al*. 2020).

### Functional enrichment analysis

Functional enrichment analysis identifies biological annotations overrepresented in either negative genetic interactions, positive genetic interactions or genetic interaction profile similarities. *P*-values are computed from the hypergeometric distribution against a background of 17,808 genes targeted by the TKOv3 CRISPR gRNA library, and false discovery rates are estimated by the Benjamini-Hochberg method. Functional annotations were obtained from Gene Ontology Biological Process, Cellular Component, and Molecular Function (Ashburner *et al*. 2000), the Molecular Signatures Database (MSigDB)(Liberzon *et al*. 2011), the EMBL-EBI Complex Portal (Meldal *et al*. 2019), and the Reactome Pathway Database (Croft *et al*. 2014; Gillespie *et al*. 2022).

### Correlation decomposition analysis

Profile similarity measures the relationship of quantitative genetic interaction (qGI) profiles between two library genes using the Pearson correlation coefficient (Billmann *et al*. 2025b). A single correlation coefficient can obscure the influence of individual query gene interactions; therefore, a correlation decomposition analysis was included to quantify the contribution of each query gene to the magnitude and direction of the overall correlation between the genetic interaction profiles associated with a pair of library genes. In this analysis, the Pearson correlation is decomposed into standardized products of deviations per screen, providing a contribution value for each screen that captures both coherent and opposing interactions between the two profiles:

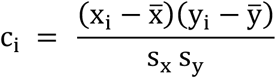

Here, *x*_*i*_ and *y*_*i*_ are the qGI scores for the two profiles in screen 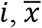 and 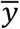 are the mean qGI scores across all screens, and *s*_*x*_ and *s*_*y*_ are the corresponding standard deviations. Outlier screens that disproportionately drive or oppose the Pearson correlation are identified using the 1.5×IQR rule.

### SAFE and RISK analysis

Spatial Analysis of Functional Enrichment (SAFE)(Baryshnikova 2016) was used to identify specific regions of the HAP1 genetic interaction profile similarity network enriched for a particular set of input genes defined by a user. A modified version of the original SAFE method was implemented for this purpose. The code and details of the modified SAFE method are available (https://doi.org/10.5281/zenodo.15320010) and described elsewhere (Billmann *et al*. 2025b).

Regional Inference of Significant Kinships (RISK)(Horecka and Rost 2026) was used as an alternative method to identify specific regions of the HAP1 genetic interaction profile similarity network enriched for a particular set of input genes. The code and details of the RISK method are available and described elsewhere (Horecka and Rost 2026).

## Results and Discussion

TheCellMap.org was originally developed to facilitate access to the global yeast genetic interaction network (Fig. 1A)(Usaj *et al*. 2017). We have now expanded the database to enable access and interactive analysis of a recently described genetic interaction network mapped in a human HAP1 cell line (Fig. 1A-B)(Billmann *et al. In press*)(Billmann *et al*. 2025b).

**Figure 1.**
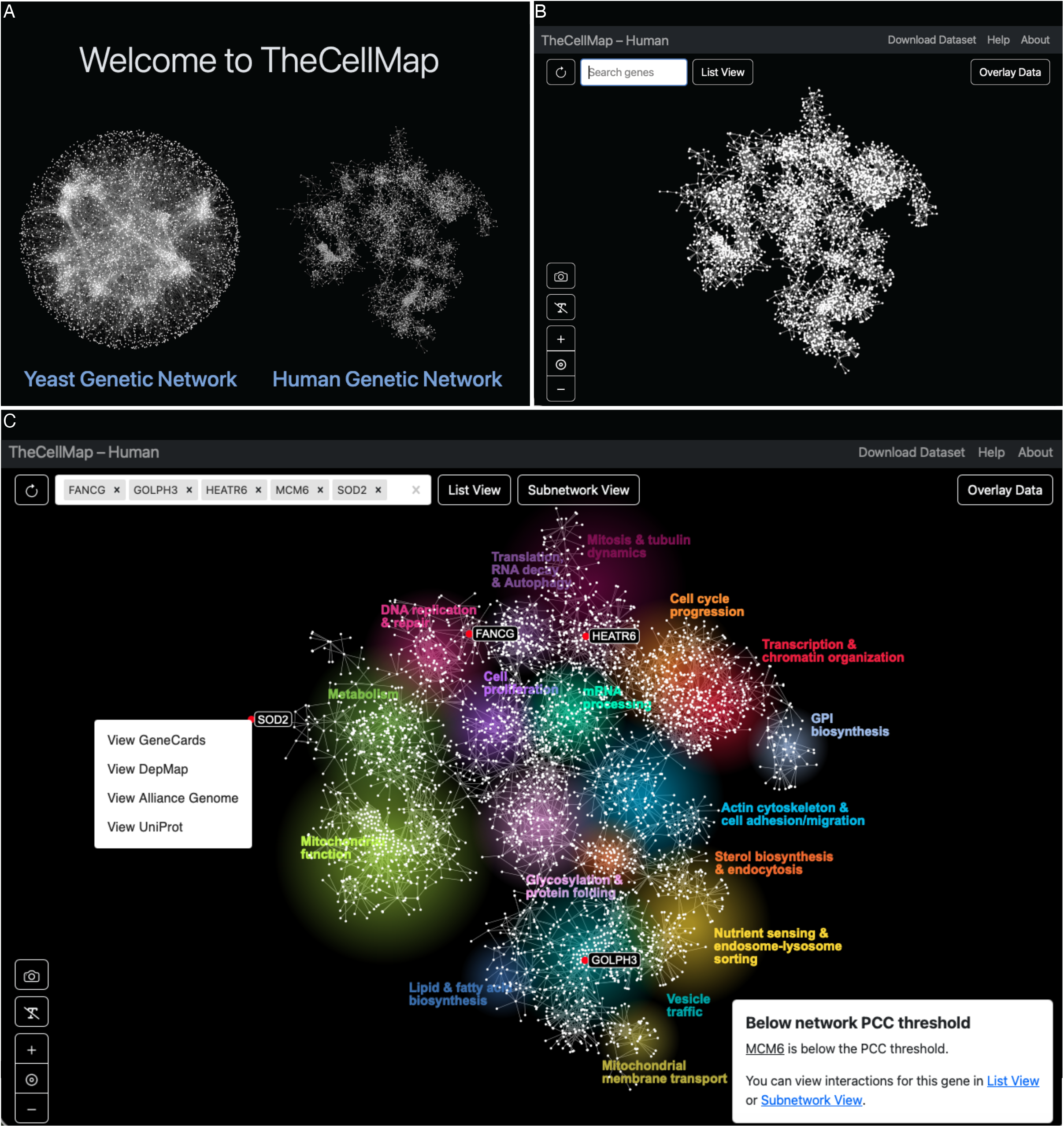
TheCellMap.org homepage. **(A)** Screenshot of TheCellMap.org homepage, where users can choose to explore genetic interactions mapped in yeast, *S. cerevisiae* (left) or human, HAP1 cell line (right). (**B**) The human HAP1 genetic interaction profile similarity network. The complete HAP1 genetic interaction dataset can be downloaded from the navigation bar. (**C**) Features of the HAP1 global network interface. Genes entered into the search window are listed in a text box and highlighted by red nodes on the network. Network clusters enriched for unique biological processes are also indicated (colored regions). Example searches show *FANCG* localized to a region of the HAP1 genetic interaction profile similarity network enriched for “DNA replication and repair”, *GOLPH3* to “Vesicle Traffic” enriched network region, and the previously uncharacterized gene, *HEATR6*, to the “Mitosis and Tubulin Dynamics” region of the HAP1 profile similarity network. Subnetworks containing selected genes can be extracted from the global network by clicking “Subnetwork View”, while profile similarities and genetic interactions can also be examined in tabular format by using “List View”. The “Overlay Data” function enables SAFE (Baryshnikova 2016) or RISK (Horecka and Rost 2026) analyses to identify regions of the HAP1 network enriched for user-defined sets of genes. Icons in the lower right allow a user to zoom in/out, hide or show gene labels, recenter the network or export the network as a PNG file. (i) Right-clicking on a node (*SOD2* shown) opens a context menu that directs the user to the corresponding Gene Page on the GeneCards Database (genecards.org), Cancer Dependency Map (DepMap) database (depmap.org), Alliance of Genome Resources (alliancegenome.org) or UniProt (uniprot.org). (ii) A warning message indicates when profile similarities for selected gene(s) do not exceed the pre-defined Pearson Correlation Coefficient threshold (PCC > 0.41) and thus are not displayed on the global HAP1 network. Hyperlinks in the warning message redirect users to corresponding subnetworks or data tables.

### HAP1 genetic interaction data acquisition, processing and scoring

A genetic interaction between two genes is quantified by comparing single mutant phenotypes, the expected double mutant phenotype, and the observed double mutant phenotype (Costanzo *et al*. 2019). For cell fitness measurements, the expected double mutant phenotype is typically modeled as a multiplicative product of the single mutant phenotypes, and genetic interactions are measured by the degree to which the observed double mutant phenotype deviates from this multiplicative expectation (Baryshnikova *et al*. 2010).

In genome-wide genetic interaction screens in human HAP1 cells, both wild-type HAP1 cells and HAP1 “query” mutant cell lines, which carry a single stable loss-of-function (LOF) mutation in a gene of interest, are transduced with the lentiviral TKOv3 gRNA library that targets ~17,000 human protein coding genes with 4-5 gRNAs/gene (Aregger *et al*. 2020; Varland *et al*. 2023; Billmann *et al*. 2025b). Genetic interactions are identified using a quantitative scoring method (qGI score) that compares the relative change in abundance of gRNAs targeting “library” genes in a query mutant background to the corresponding changes measured across 39 genome-wide screens performed with the parental wild-type (WT) HAP1 cell line (Billmann *et al*. 2025a). Negative genetic interactions correspond to library genes whose gRNAs show significantly decreased abundance in a query mutant background relative to WT, whereas positive genetic interactions correspond to library genes with increased gRNA abundance in a query mutant cell line relative to WT (Billmann *et al*. 2025a; Billmann *et al*. 2025b). Notably, the qGI score provides a quantitative indicator of functional relatedness, as stronger negative interaction scores were associated with gene pairs that shared closer functional relationships (Billmann *et al*. 2025b). To evaluate reproducibility and estimate false negative and false positive rates across a range of qGI score and statistical confidence thresholds, screens for 50 query mutant cell lines were repeated multiple times (Billmann *et al*. 2025a; Billmann *et al*. 2025b). In total, we performed ~300 genome-wide CRISPR-Cas9 knockout screens using 222 unique query mutant cell lines to analyze ~4 million double mutants and construct a HAP1 genetic network comprising ~89,000 quantitative genetic interactions (Billmann *et al*. 2025b).

The HAP1 genetic interaction network includes ~47,000 negative and ~42,000 positive genetic interactions (Billmann *et al*. 2025b). Genetic interaction profile similarities were also calculated by computing Pearson correlation coefficients (PCC) between all pairs of library genes targeted by the TKOv3 gRNA library (Billmann *et al*. 2025b). The complete unfiltered dataset, including negative and positive genetic interactions as well as genetic interaction profile similarities, is available for download in various formats from TheCellMap.org navigation bar (Fig. 1B).

TheCellMap.org also enables users to explore and visualize genetic interaction data for selected genes in multiple formats, including: (1) a global genetic interaction profile similarity network for selected genes (“Subnetwork View”) and (2) lists of gene pairs ranked based on either genetic interaction profile similarity or the magnitude of negative or positive qGI scores (“List View”)(Fig. 1C). Because the genetic interaction profile similarity network provides a resource for annotating gene function (Costanzo *et al*. 2010; Costanzo *et al*. 2016; Billmann *et al*. 2025b), we implemented an “Overlay Data” tool that also allows users to test whether specific regions of the HAP1 genetic interaction profile similarity network are statistically enriched for a user-defined gene set (Fig. 1C). All four data visualization modes are described in detail below.

### Navigating the global genetic interaction profile similarity network

TheCellMap.org provides an interactive visualization of the HAP1 genetic interaction profile similarity network (Fig. 1B-C). In this network, nodes represent library genes and edges connect library gene pairs that share similar patterns of genetic interactions, defined by genetic interaction profile similarities exceeding a Pearson Correlation Coefficient (PCC) threshold of > 0.41. As described previously (Billmann *et al*. 2025b), the complete HAP1 genetic interaction dataset underwent a series of normalization procedures to remove non-specific interaction signals prior to computing profile similarities thus generating a high confidence network encompassing ~3,600 human genes.

To visualize the network, a layout algorithm was applied to position genes (nodes) in two-dimensional space, such that genes with more similar genetic interaction profiles (i.e. higher PCC values) are placed closer together, whereas genes with less similar genetic interaction profiles are placed farther apart (Costanzo *et al*. 2016). Compared to the global yeast profile similarity network, the HAP1 network is relatively sparse because each library gene profile is derived from genetic interactions with only 222 unique query genes. In total, the current HAP1 genetic interaction profile similarity network consists of 3,784 library genes connected by 6,433 edges. Approximately 74% (2787/3784) of the genes reside within large, visually discernible network clusters (Fig. 1C)(Billmann *et al*. 2025b). As observed in the global yeast network (Costanzo *et al*. 2016), densely connected clusters in the HAP1 profile similarity network are enriched for functionally related genes within distinct biological processes (e.g. Vesicle Traffic, GPI Biosynthesis etc.). Each bioprocess can be further resolved into more refined gene clusters representing specific pathways and protein complexes (Billmann *et al*. 2025b).

The specific location of one or more genes can be mapped onto the global profile similarity network by entering the gene name(s) into the search window (Fig. 1C). After a search, the selected gene(s) are highlighted on the network as red node(s) (Fig. 1C). The corresponding Gene Page on the GeneCards Database (genecards.org), Cancer Dependency Map (DepMap) database (depmap.org), Alliance of Genome Resources (alliancegenome.org) or UniProt (uniprot.org) are available by right-clicking on a selected gene (Fig. 1C, box labelled “I”). Functions located in the bottom left include a camera icon that allows users to export the currently displayed network as a Portable Network Graphics (.png) image along with buttons to zoom in/out, re-center the network and hide gene labels (Fig. 1C). Following a gene search, the major biological processes associated with each network cluster, as determined by Spatial Analysis of Functional Enrichment (SAFE)(Baryshnikova 2016), are also highlighted using distinct colors (Fig. 1C). SAFE identifies dense network regions associated with specific functional attributes. Applying SAFE to the HAP1 genetic interaction profile similarity network revealed 427 significantly enriched Gene Ontology (GO) bioprocess terms (Ashburner *et al*. 2000) that mapped to 17 unique bioprocess-level network regions encompassing 2,787 genes (Billmann *et al*. 2025b).

The global genetic interaction profile similarity network provides a powerful resource for predicting gene function because the network position of a given gene often reflects its biological role (Costanzo *et al*. 2010; Costanzo *et al*. 2016; Billmann *et al*. 2025b). For example, *FANCG*, a gene involved in DNA recombination (Garcia-Higuera *et al*. 2001), localizes to the “DNA replication and repair” bioprocess-enriched cluster, consistent with its established function. *GOLPH3*, which encodes a phosphatidylinositol-4-phosphate-binding protein involved in Golgi membrane trafficking (Dippold *et al*. 2009), maps to the “Vesicle Traffic” region of the HAP1 profile similarity network (Fig. 1C). A previously uncharacterized gene, *HEATR6*, resides within a bioprocess cluster enriched for genes involved in Mitosis and tubulin dynamics, suggesting that *HEATR6* may share a functional role with other genes involved in these functions (Fig. 1C). A new gene search can be initiated by clicking on the “Reset Graph” icon in the upper left corner. Clicking “TheCellMap” logo in the navigation bar redirects to the home page where a user can select between yeast and human genetic interaction datasets (Fig. 1A, C).

As described above, a similarity threshold was applied to the genetic interaction profile similarity matrix to reduce complexity of the complete HAP1 network and facilitate visualization of the biological process-enriched clusters (Fig. 1C)(Billmann *et al*. 2025b). Consequently, any library gene whose genetic interaction profile does not exceed the minimum similarity threshold (PCC > 0.41) with at least one other library gene, such as *MCM6* (Fig. 1C), is not displayed on the global network. Searching for such genes triggers a warning message indicating that the selected gene is not represented on the global network (Fig. 1C, box labelled “ii”). However, interaction data for these genes can be accessed either as a subnetwork or as a ranked list of profile similarities by clicking the hyperlink provided in the warning message box (Fig. 1C, box ii) or by clicking the corresponding buttons next to the search window. Features associated with “Subnetwork View” and “List View” formats are described in detail below.

### Extracting genetic interaction profile similarity subnetworks

Once a gene or set of genes is selected, the connections that define its position on the HAP1 genetic interaction profile similarity network can be examined in greater detail by clicking the “Subnetwork View” button (Fig. 1C). The “Subnetwork View” displays the gene(s) of interest together with all library genes directly connected to it, as well as neighboring genes that are indirectly connected through one additional gene/node, thereby revealing distinct subnetwork clusters (Fig. 2). Subnetwork edges connect genes with similar genetic interaction profiles and are measured as PCCs. Because these genes are located close to the gene of interest on the HAP1 profile similarity network, they must share similar sets of genetic interactions with the gene of interest. The same network layout algorithm used for the global network is applied to reposition nodes based solely on the connections present within the subnetwork (Fig. 2). Node color reflects the biological processes assigned in the global HAP1 network, as determined by SAFE analysis (Figs. 1C, 2). For example, as noted above, most genes within the *FANCG* or the *GOLPH3* profile similarity subnetworks map to the DNA replication and repair and Vesicle Traffic, respectively (Fig. 2A-B). The *HEATR6* profile similarity subnetwork is primarily composed of genes involved in Mitosis and tubulin dynamics (Fig. 2C). All functions available when right-clicking a node in the global HAP1 network (Fig. 1C) are also available within the Subnetwork View.

**Figure 2.**
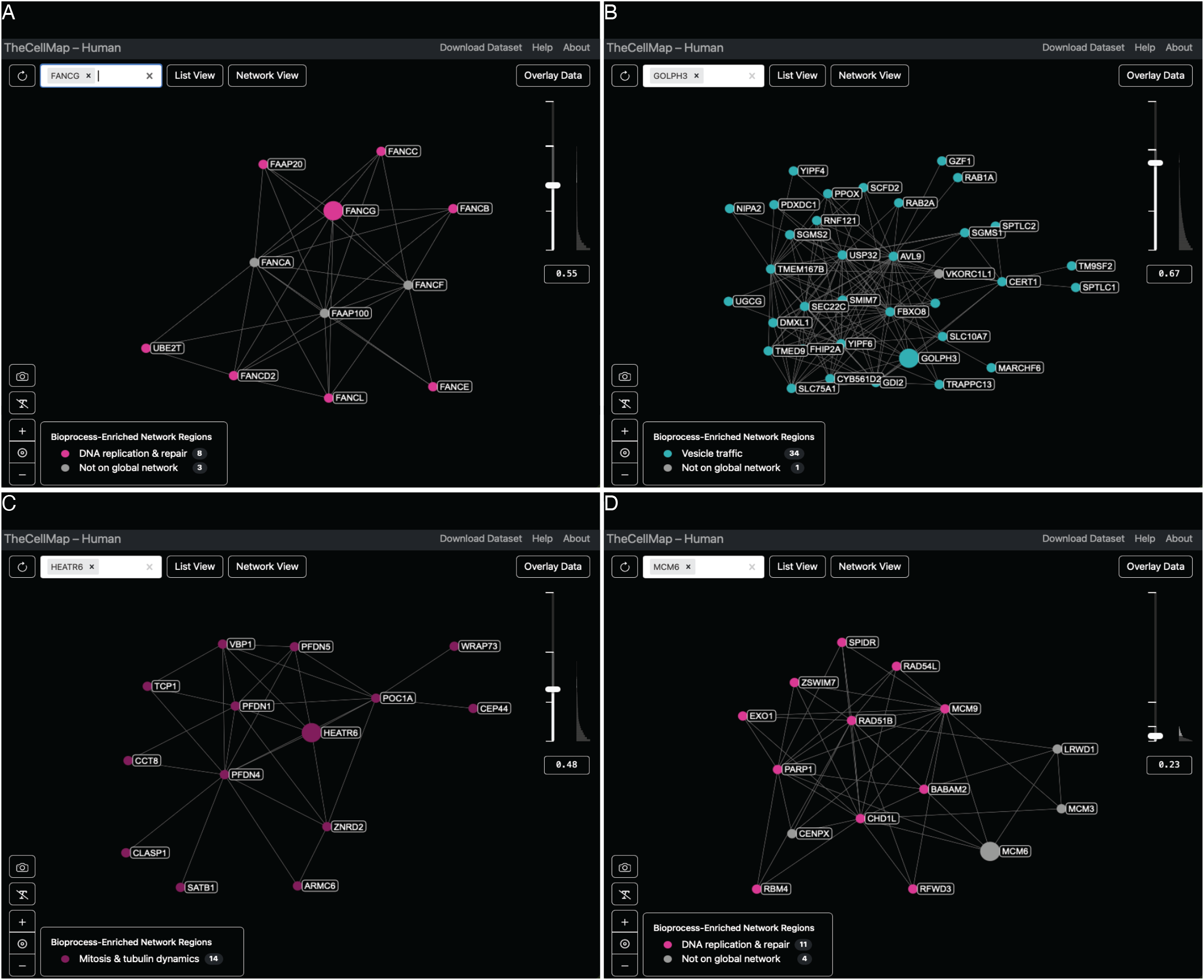
Genetic interaction profile similarity subnetwork view. (**A**) *FANCG*-specific profile similarity sub-network generated using a user-defined PCC threshold (PCC > 0.55). Genes with genetic interaction profiles similar to *FANCG* are displayed and positioned in an unbiased layout based on the extent of similarity to the *FANCG* profile. Nodes are colored according to their annotation in the global HAP1 network. (**B**) *GOLPH3*-specific profile similarity sub-network generated using a user-defined PCC threshold (PCC > 0.67). Nodes are colored according to their annotation in the global HAP1 network. (**C**) *HEATR6*-specific profile similarity sub-network generated using a user-defined PCC threshold (PCC > 0.48). Nodes are colored according to their annotation in the global HAP1 network. (**D**) *MCM6*-specific genetic interaction profile similarity subnetwork generated using a more lenient user-defined profile similarity threshold (PCC > 0.23). Nodes are colored according to their annotation in the global HAP1 network.

A slider bar allows users to highlight profile similarities above or below a specified PCC threshold and is accompanied by a histogram that displays the distribution of edge density across threshold values. The default threshold for each subnetwork is dynamically determined based on the selected gene(s). Specifically, the threshold is chosen to minimize the difference between the number of edges connected to the gene of interest and the log2 of the number of nodes in the resulting subnetwork (Fig. 2A-C). Once the subnetwork is displayed, users may manually adjust the similarity threshold using the slider bar or by directly inputting a PCC threshold value (Fig. 2D). When the threshold is changed, the same layout algorithm is automatically applied to re-organize the network according to nodes and edges that satisfy the new PCC threshold. For example, *MCM6*, which encodes a component of the pre-initiation complex required for initiation of DNA replication (Takisawa *et al*. 2000), does not appear on the global network because its strongest genetic interaction profile similarity does not exceed the global network PCC threshold (PCC > 0.41, Fig. 1C). However, by applying a more lenient but significant profile similarity threshold (e.g. PCC > 0.2), a genetic interaction profile similarity subnetwork emerges that includes *MCM6* along with other genes involved in DNA damage and repair (Fig. 2D). The minimum, maximum and global profile genetic interaction similarity network PCC thresholds are displayed adjacent to the slider bar.

### Exploring genetic interaction profile similarities in List View

In addition to visualizing HAP1 genetic interaction profiles as networks, profile similarities for a selected gene or set of genes can also be viewed in a tabular format (Fig. 3). The table format is accessed by clicking the “List View” button from either the global network page or the subnetwork page (Fig. 1C). The List View interface includes two menus: “Data Type” and “Selected Genes” (Fig. 3A). When Profile Similarity is selected from the Data Type menu, the library gene highlighted in the Selected Genes menu appears in the “Library Gene A” column of the main table, and other library genes that share profile similarities with the selected gene are displayed in the “Library Gene B” column, ordered in descending rank according to their genetic interaction profile similarity (PCC) (Fig. 3A). For example, selecting *FANCG* as Library Gene A produces a ranked list of Library Gene B entries sorted by similarity to the *FANCG* library gene interaction profile (Fig. 3A). Consistent with prior observations that genes in the same pathway tend to share similar genetic interaction profiles (Costanzo *et al*. 2019), *FANCG* shows its strongest similarity to *FANCA*, whose gene product functions in the same DNA recombination pathway (Fig. 3A). By default, only genes with genetic interaction profiles exceeding a minimum similarity threshold (PCC > 0.1) with the selected Library Gene A are included in the Library Gene B column. The table displays data for the gene highlighted in the Selected Genes menu. In addition, hovering over any row in the profile similarity table reveals icons linking to external resources, including GeneCards (genecards.org), DepMap Database (depmap.org) and the Alliance of Genome Resources (alliancegenome.org), allowing users to quickly access detailed gene-specific information (Fig. 3A).

**Figure 3.**
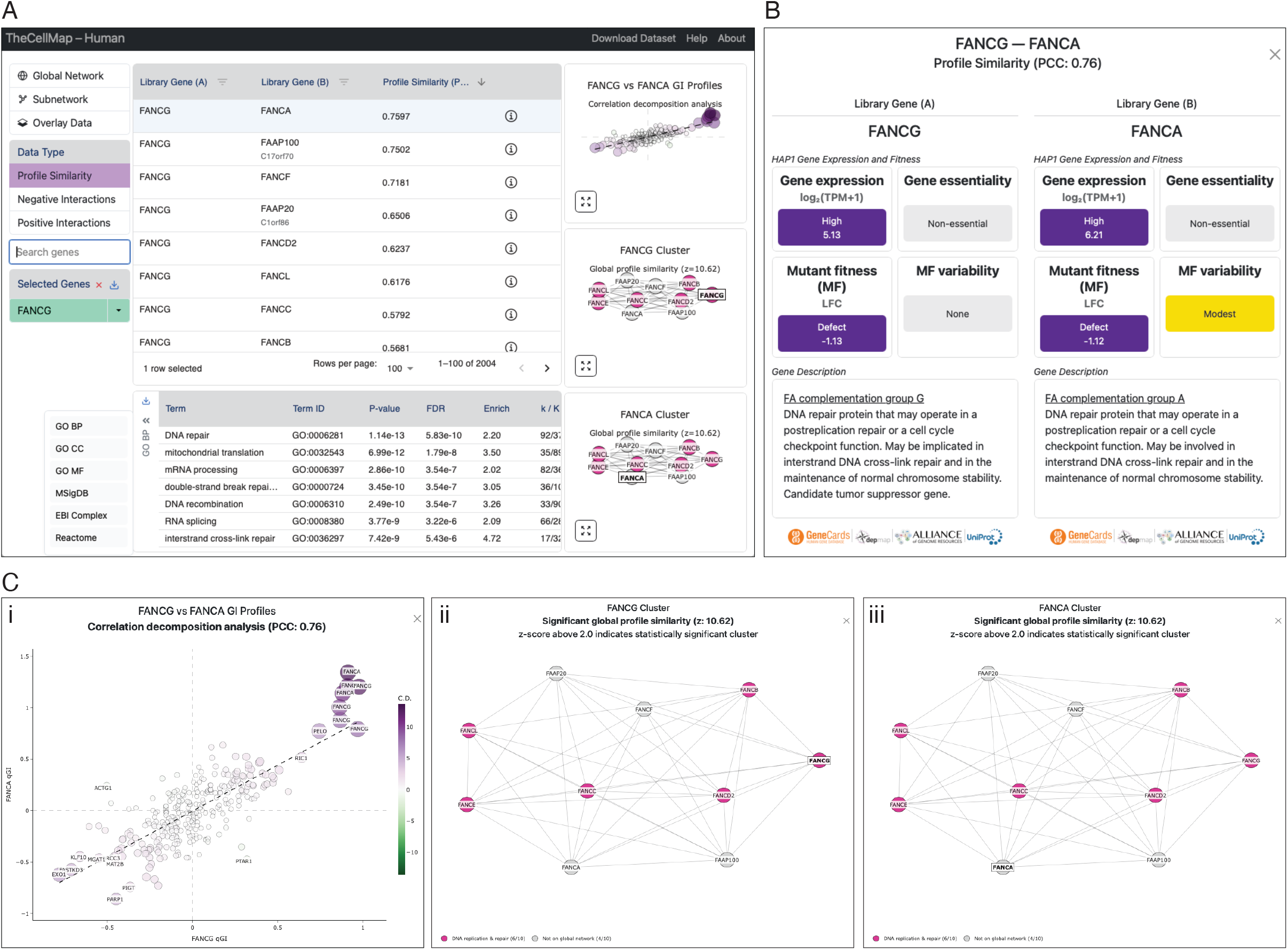
Profile similarities visualized in List view. **(A)** Screenshot illustrating genetic interaction profile similarities for a *FANCG* query gene displayed in tabular format. From the “Data Type” menu, users can select genetic interaction profile similarity data. The searched gene is shown in the “Selected Genes” panel. A dropdown menu associated with each selected gene allows users to modify the gene list or access additional gene-specific resources. The main table lists profile similarities for the gene highlighted in the Selected Genes panel. Links to external gene resources appear below genes in the selected column. Clicking the information icon at the far right of the selected row in the main table reveals additional information pertaining to the gene pair (see panel B). Previews of the correlation decomposition scatter plot and library gene cluster visualizations for the gene pair in the selected row of the main table. Each graph, described in (C), can be enlarged using the expand icon in the lower left corner. A table summarizing results from automated functional enrichment of genes (Library Gene B column of main table) that share similar genetic interaction profiles with the highlighted gene in the Selected Genes panel. By default, enrichment is tested for GO biological process terms. Alternative functional annotation standards can be selected from the dropdown menu. **(B)** Screenshot of the gene information window, which displays HAP1 gene expression and mutant fitness phenotypes for a specific gene pair (e.g. *FANCG-FANCA*) selected in the main table (see panel A). For expression and mutant fitness (MF), dark purple indicates high expression and/or a strong single mutant fitness defect in HAP1 cells, light purple indicates average expression and/or fitness, and grey indicates no detectable fitness phenotype. For MF variability, genes with highly variable (orange) or modestly variable (yellow) single mutant fitness phenotypes are indicated, whereas genes with stable single mutant fitness across wild type control screens are shown in grey. **(C)** (i) Enlarged views of the correlation decomposition scatter plot. Scatter plot showing genetic interactions (qGI scores) for *FANCG* (x-axis) and *FANCA* (y-axis) library genes across 298 query genes (nodes). Node color reflects correlation decomposition scores. Purple nodes indicate positive contributions to profile similarity, where *FANCG* and *FANCA* library genes show similar genetic interactions with the same query genes, whereas green nodes indicate negative contributions to profile similarity where *FANCG* and *FANCA* show opposite genetic interactions with the same query gene. Node size is proportional to the absolute value of the correlation decomposition score. (ii-iii) Enlarged views of library gene clusters corresponding to the gene pair highlighted in the main table derived from hierarchical clustering of the full genetic interaction dataset, as described in (Billmann *et al*. 2025b). Node color corresponds to functionally enriched network regions shown in Fig. 1C. Grey nodes indicate genes not included or annotated in the global network. Although *FANCG* and *FANCA* belong to the same cluster in this example, genes can belong to distinct but functionally related clusters (Billmann *et al*. 2025b).

Each row in the profile similarity table can also be expanded by clicking on the information icon at the far right of the table, revealing additional details about the highlighted gene pair (Fig. 3A-B). This expanded view includes functional descriptions, gene expression level and essentiality status (essential vs. nonessential) in HAP1 cells, as well as single mutant fitness measurements derived from a panel of genome-wide CRISPR-Cas9 knockout screens in a WT HAP1 cell line (Billmann *et al*. 2025a; Billmann *et al*. 2025b)(Fig. 3B). In particular, we previously showed that a small set of library genes exhibit highly or moderately variable single mutant fitness effects across replicate genome-wide WT screens (e.g. *FANCA*)(Billmann *et al*. 2025a; Billmann *et al*. 2025b). The gene information page also indicates if a selected gene pair involves a gene associated with variable mutant fitness along with other gene-specific links to external resources (Fig. 3B).

Data for all genes listed in the Selected Genes menu can be downloaded by clicking the “Download” icon in the Selected Genes menu header (Fig. 3A). Alternatively, data for individual genes can be downloaded using the dropdown menu associated with each selected gene. Files are exported in a Microsoft Excel (.xlsx) format and include all three data types: Profile Similarity, Negative Interactions and Positive Interactions for the selected gene(s). The dropdown menu for each gene also provides options to remove that gene from the Selected Genes list or to navigate directly to its Gene Page on different external databases.

Each gene pair (i.e. row) in the profile similarity table is accompanied by three preview graphs, arranged vertically to the right of the table (Fig. 3A). Preview graphs for a gene pair are displayed when the corresponding row is selected and each preview graph can be expanded by clicking the icon in the bottom left of the preview window (Fig. 3A, 3C). Gene labels become visible when hovering over individual nodes in either the preview or the expanded graph windows. Users can zoom in/out, select specific regions of the plot or save the visualization as a .png file from the expanded graph window view.

### Correlation decomposition analysis of genetic interaction profiles

Profile similarity measures the relationship of genetic interaction profiles between two library genes using PCCs (Billmann *et al*. 2025b). To identify specific genetic interactions that drive similarity between genetic interaction profiles of two library genes, we implemented a correlation decomposition approach, which quantifies the contribution of each individual genetic interaction to the magnitude and direction of the overall correlation between the genetic interaction profiles associated with a given pair of library genes (Methods). A graph depicting the correlation decomposition of genetic interaction profiles for a selected library gene pair is shown in the preview window in the top right of the Profile Similarity List View page (Fig. 3A). Nodes in the plot represent 298 genome-wide genetic interaction screens performed with 222 unique HAP1 query mutant cell lines, described previously (Billmann *et al*. 2025b). Genetic interaction scores (qGI scores) measured between the set of 298 query genes and library gene A are plotted on the x-axis while the qGI score associated with an interaction between the 298 query genes and library gene B are plotted on the y-axis (Fig. 3C i). Purple nodes indicate query genes whose interactions with both library genes are similar in sign and magnitude, thereby contributing strongly and positively to the overall profile similarity between library genes A and B. For example, we carried out a *PELO* query gene screen, and it identified positive qGI scores for both the *FANCA* and *FANCG* library genes, such that these shared interactions contribute to the positive correlation for these two library genes. Two independent *FANCA* query gene screens identified positive interactions with *FANCA* (i.e. self-interaction)(Billmann *et al*. 2025b) and *FANCG* library genes, which reinforces that finding that nonessential genes belonging to the same pathway or complex tend to be connected by positive interactions, and these shared interactions contribute to the positive correlation for these two library genes (**Fig. 3C i**)(Costanzo *et al*. 2019; Billmann *et al*. 2025b). The *EXO1* query gene showed negative qGI scores for both *FANCA* and *FANG* library genes, and these shared interactions further contribute to the correlated profile similarity (**Fig. 3C i**). In contrast, green nodes, such as *PTAR1*, represent query genes that show opposing qGI scores with library gene A and library gene B (e.g. a positive qGI score with *FANCG* and a negative qGI score with *FANCA*), resulting in negative correlation decomposition (**Fig. 3C i**).

### Genetic interaction clusters

To complement the global HAP1 genetic interaction profile similarity network, we applied a hierarchical clustering algorithm to the complete HAP1 genetic interaction dataset. This analysis identified ~4,800 clusters or modules of library genes with similar genetic interaction profiles, a subset of which correspond to genes that function together in the same bioprocess, pathway or protein complex (Billmann *et al*. 2025b). When a user selects a row in the Profile Similarity table, two additional preview graphs appear below the correlation decomposition scatter plot (Fig. 3A). These graphs display the hierarchical clustering modules that contain the corresponding Library Genes A and B, respectively (Fig. 3A, 3C ii-iii). The statistical significance of each module is provided in the expanded graph window (Fig. 3C ii-iii). A z-score greater than 2.0 indicates that the average profile similarity among genes in the module is significantly higher than expected by chance (Billmann *et al*. 2025b). Nodes are colored according to the bioprocess enriched region to which they map on the global HAP1 profile similarity network (Fig. 1C). Because hierarchical clustering was performed on the full, unfiltered genetic interaction dataset, modules derived from this approach include genes that do not appear on the global HAP1 profile similarity network (Fig. 1)(Billmann *et al*. 2025b). These genes are shown as grey nodes in the cluster plots (Fig. 3C ii-iii). For example, *FANCG* clusters with other genes in the Fanconi Anemia complementation group including, *FANCA* and *FANCF*, which are not represented on the HAP1 global profile similarity network but belong to the same hierarchical cluster module (Fig. 3C ii-iii)(Billmann *et al*. 2025b). Most significant gene clusters (~93%, 384/412) were enriched for specific GO bioprocess terms that span diverse cellular functions (Billmann *et al*. 2025b).

### Automated functional enrichment analysis

TheCellMap.org enables automated functional enrichment analysis for gene lists derived from profile similarity, negative genetic, and positive genetic interaction data (Fig. 3A). When Profile Similarity is selected from the Data Type menu, all genes in the Library Gene (B) column whose similarity to the selected Library Gene (A) exceeds PCC > 0.1 are automatically tested for enrichment using a hypergeometric test. For negative and positive genetic interaction Data Types, enrichment is performed on the column opposite the searched gene; if the searched gene is a query, genes in the Library Gene column are tested and vice versa (Methods).

Gene Ontology Biological Process (GO BP) terms are used as the default annotation standard, and enrichment results are displayed in a dedicated table at the bottom of the page (Fig. 3A). For example, library genes with profile similarity to *FANCG* (PCC > 0.1) are enriched for terms related to DNA repair (Fig. 3A). As described previously (Rahman *et al*. 2021), genes involved in mitochondrial-related functions are among the most highly connected and highly correlated genes in the HAP1 genetic interaction profile similarity network. Thus, HAP1 interaction profiles, as also observed for DepMap co-essentiality profiles, often show statistical enrichment for mitochondrial-related functional terms (Fig. 3A)(Rahman *et al*. 2021). Evidence suggests that the elevated interaction frequency associated with mitochondrial-related genes may be related to the increased stability of proteins encoded by these genes, which can impact detection of mutant growth phenotype until the corresponding WT protein is depleted in the context of a pooled genome-wide CRISPR screen. As a result, genetic interaction profiles over-represented for mitochondrial-related genes should be explored with caution (Rahman *et al*. 2021; Billmann *et al*. 2025b).

In addition to GO bioprocess, other functional standards including GO Cellular component (GO CC), GO Molecular Function (GO MF), Molecular Signatures Database (MSigDB), EBI Protein complex portal (EBI Complex) and Reactome Pathways, can be selected for enrichment analysis from the dropdown menu (Fig. 3A).

### Exploring negative and positive genetic interactions

TheCellMap.org also enables detailed exploration of negative and positive genetic interactions for query and/or library genes of interest. These interactions can be explored in tabular format by clicking the “List View” button for a selected gene(s) on the global network page (Fig. 1C). When “Negative Interaction” is chosen from the Data Type menu (Fig. 4A), all negative genetic interactions involving the highlighted gene in the Selected Gene menu (Fig. 4A) are displayed in the interaction table (Fig. 4A). Positive interactions for the selected gene can be accessed by choosing “Positive Interactions” from the Data Type menu (Fig. 4B). Using the dropdown menu next to any gene listed in the Selected Genes menu, users may choose to view “All Interactions” for that gene, aggregating results from when it was screened as a query and/or library gene, or restrict the display to only the negative or positive interactions identified when the gene was screened specifically as a query or specifically as a library gene (Fig. 4A-B). For query genes that were screened multiple times, users may further choose to retrieve the complete set of negative or positive interactions from all biological replicate screens or limit the view to those interactions obtained from a single screen associated with a unique identifier number (GIN ID) (Fig. 4A). Only data corresponding to the gene currently highlighted in the Selected Genes menu are displayed in the table.

**Figure 4.**
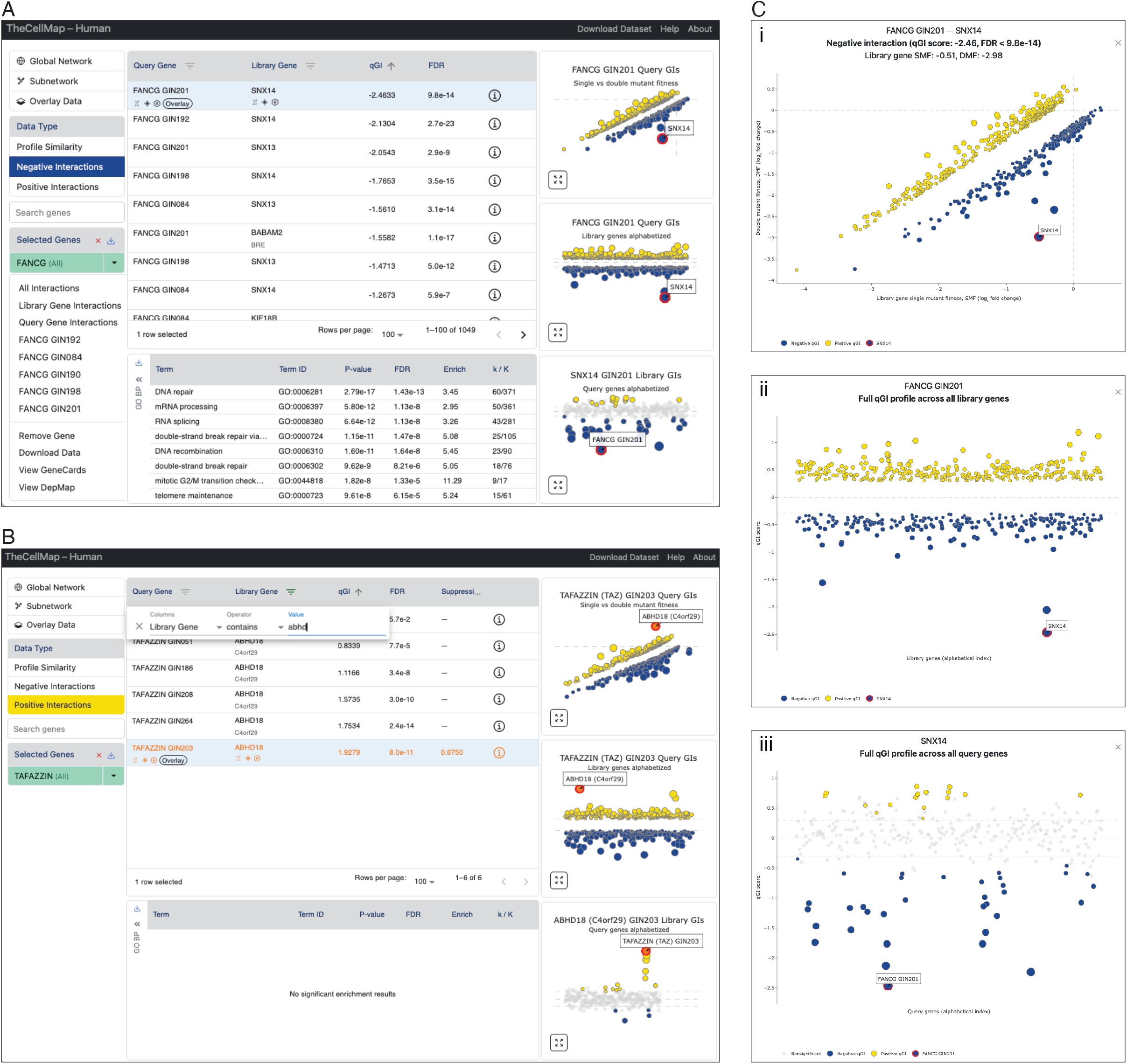
Genetic interaction table. **(A)** Screenshots illustrating negative genetic interactions for the indicated query genes displayed in a tabular format. From the “Data Type” menu, users can select to view negative genetic interaction data. The highlighted gene is shown in the “Selected Genes” panel. A dropdown menu associated with each gene allows a user to modify the selected gene list, by viewing interactions identified for a gene as a library and/or query gene, examine data from individual replicate screens or navigate to additional gene-specific resources. The main data table lists genetic interactions for a gene highlighted in the Selected gene menu. Links to external gene resources and Data Overlay analysis appear below genes in the selected column. Clicking the information icon reveals additional information for the corresponding gene pair (see Fig. 3B). Previews of genetic interaction scatter plots for gene pairs shown in the selected row of the main table. Each plot, described in (C), can be enlarged using the expand icon in the lower left corner. A table summarizing results from automated functional enrichment analysis of genes that exhibit genetic interactions with the selected gene. By default, enrichment is performed using GO biological process terms. Alternative functional annotation standards can be selected from the dropdown menu. (**B**) Screenshots illustrating negative genetic interactions for the indicated query genes displayed in a tabular format. The same functions described for negative interactions, highlighted in panel A, are also available from the positive interaction List View. Genetic interactions can be filtered using the filter icons next to the Query or Library gene headers to display genetic interactions involving specific gene pairs. Strong positive interactions consistent with potential suppression interactions (Suppressor score > 0.5)(Billmann *et al*. 2025b), such as *TAFAZZIN-ABHD18*, are indicated in orange text. (**C**) Enlarged views of genetic interaction scatter plots for a selected query gene or a selected library gene corresponding to the gene pair highlighted in the main table (i.e. table row). (i) Scatterplots depicting genetic interactions for the indicated query gene, with library gene single mutant fitness (from loss-of-function screens in a WT HAP1 cell line) plotted on the x-axis and the corresponding double mutant fitness in a HAP1 cell line carrying an additional loss-of-function mutation in the query gene plotted on the y-axis (Billmann *et al*. 2025a; Billmann *et al*. 2025b). Negative (blue) and positive (yellow) genetic interactions that satisfied a standard genetic interaction threshold (|qGI| > 0.3, FDR < 0.1) are shown. (ii) Negative (blue) and positive (yellow) genetic interactions that satisfied a standard genetic interaction threshold (y-axis, |qGI| > 0.3, FDR < 0.1) for the indicated query gene, plotted in alphabetical order of tested library genes (x-axis). (iii) Negative (blue) and positive (yellow) genetic interactions that satisfied a standard genetic interaction threshold (y-axis, |qGI| > 0.3, FDR < 0.1) for the indicated library gene, plotted in alphabetical order of tested query genes (x-axis). In all panels, nodes outlined in red denote the specific genetic interaction corresponding to the highlighted row in the main table.

The “Query Gene” column in the Negative or Positive Interaction table indicates the HAP1 cell line carrying a stable LOF mutation in the specified query gene (Fig. 4A-B). The query gene name is followed by a unique screen identification number (GIN ID number), which distinguishes independent biological replicate screens using the same query mutant cell line (e.g. *FANCG* query gene screen GIN201 vs. GIN192). The Library Gene column lists the gene targeted by the TKOv3 gRNA library that shows a negative or positive interaction with the indicated query gene (Fig. 4A-B). For example, in genome-wide screen FANCG GIN201, a HAP1 cell line carrying a stable LOF mutation in *FANCG* showed a negative interaction with *SNX14*, which encodes a sorting nexin involved in vesicle trafficking (Fig. 4A), while screen TAFAZZIN GIN203 identified a positive interaction between the *TAFAZZIN* query gene, involved in cardiolipin biosynthesis, and the *ABHD18* library gene, which encodes a newly discovered lipase enzyme that functions upstream of TAFAZZIN in the cardiolipin biosynthetic pathway (Fig. 4B)(Masud *et al*. 2025). The qGI score compares the abundance of gRNAs derived from CRISPR-based screens in WT HAP1 cells, which reflects library gene single mutant fitness, to the abundance of the same TKOv3 gRNAs in HAP1 query mutant cells carrying a stable mutation in a query gene of interest, which provides an estimate of double mutant fitness (Billmann *et al*. 2025a). Negative qGI scores reflect genes with gRNAs that show significantly decreased abundance in a query mutant relative to WT (Fig. 4A). Extreme negative interactions represent synthetic lethal interactions, which tend to have stronger negative qGI scores while less extreme negative interactions, like synthetic sick interactions, have weaker negative qGI scores (Billmann *et al*. 2025b). In contrast, positive qGI scores identify genes with increased gRNA abundance in a query mutant relative to WT (Billmann *et al*. 2025a) and gene pairs with extremely strong positive qGI scores may reflect instances of genetic suppression, which are described in more detail below (Fig. 4B). The associated False Discovery Rate (FDR) column reflects the statistical confidence in the qGI score (Fig. 4A-B)(Billmann *et al*. 2025a). Only interactions that satisfy a standard confidence threshold (|qGI| > 0.3 and FDR < 0.1) are displayed in the interaction tables (Fig. 4A-B)(Billmann *et al*. 2025a; Billmann *et al*. 2025b). Similar to the Profile Similarity table (Fig. 3A), each row in the Interaction table can be expanded using the information icon at the far right (Fig. 4A), revealing function descriptions, HAP1 expression level, essentiality status, single mutant fitness and variability measurements, along with gene-specific resources (Fig. 3B). Negative and positive genetic interaction datasets can be downloaded for all genes listed in the Selected Genes menu using the “Download” icon in the menu header (Fig. 4A). Data for individual genes can also be downloaded via the dropdown menu associated with each selected gene.

Each row in the negative or positive interaction table is also accompanied by three preview graphs displayed to the right of the table (Fig. 4A-B). The graphs corresponding to a specific gene pair are shown when the row is selected (Fig. 4A-B), and each graph can be expanded by clicking on the icon in the bottom left corner of the preview window (Fig. 4C). The first graph plots library gene single mutant fitness on the x-axis and double mutant fitness for the same library gene in the selected query gene mutant background on the y-axis (Fig. 4C i). The qGI score reflects the deviation between single and double mutant fitness values. Gene pairs associated with significant negative or positive qGI scores (|qGI| > 0.3 and FDR < 0.1) are depicted as blue and yellow nodes, respectively (Fig. 4C i). Nodes outlined in red correspond to the specific genetic interaction selected in the table (Fig. 4C i). Users can further refine the interaction table by clicking the filter icon next to the “Query Gene” or “Library Gene” table headers and entering comma-separated keywords to restrict the display to interactions involving specific gene(s) of interest (Fig. 4B).

A subset of extreme positive interactions where the double mutant fitness is at least 50% greater than the fitness of the sickest single mutant, represents potential genetic suppression interactions (Billmann *et al*. 2025b). These candidate suppressor interactions are highlighted in orange in the positive interaction table and shown as orange nodes in the corresponding genetic interaction plots (Fig. 4B). For example, HAP1 genetic interaction analysis identified *ABHD18* as a previously unrecognized component of cardiolipin biosynthesis, whose inactivation suppresses Barth Syndrome disease gene (*TAFAZZIN*)-associated phenotypes (Fig. 4B)(Billmann *et al*. 2025b; Masud *et al*. 2025).

The two remaining graphs (Fig. 4C ii-iii) display the full set of negative and positive interactions identified for the specific query-library gene pair highlighted in the main table. The “Query GIs” graph plots all significant negative and positive qGI scores associated with the selected query gene on the y-axis, with all tested library genes arranged alphabetically along the x-axis (Fig. 4C ii). Conversely, the “Library GIs” graph plots all significant genetic interactions measured for the selected library gene on the y-axis, with all tested query genes arranged alphabetically on the x-axis (Fig. 4C iii). The relative sparsity of the Library GI graph reflects the structure of the HAP1 genetic interaction dataset. The network is based on 298 genome-wide screens corresponding to 222 unique query genes, whereas each Query GIs graph includes the thousands of library genes tested in every screen. The identity of genetic interactions in either plot can be revealed by hovering over individual nodes in the expanded graph window (Fig. 4C). Users can also pan, zoom in/out, select specific plot regions or save the figure as a .png file from the expanded graph window.

### Overlaying negative and positive genetic interactions onto the profile similarity network

Like the Profile Similarity table (Fig. 3A), hovering over a row in the Negative and Positive Interaction table reveals icons that link directly to the corresponding gene pages in GeneCards (genecards.org), the DepMap Database (depmap.org) or the Alliance of Genome Resources (alliancegenome.org)(Fig. 4A). By clicking the fourth icon, labeled “Overlay”, located beneath each query gene, users can perform automated functional analysis for the set of negative and positive genetic interactions associated with that query gene (Figs. 4A). By default, choosing the Overlay icon initiates a SAFE (Spatial Analysis of Functional Enrichment) analysis, which identifies regions of the HAP1 profile similarity network that are statistically overrepresented for a query gene’s set of negative and positive interactions. As an alternative to SAFE, enrichment of query gene genetic interactions can also be assessed using Regional Inference of Significant Kinships (RISK), a recently developed method that integrates clustering algorithms, rigorous statistical analysis, and a visualization framework to enhance interpretation of biological networks (Horecka and Rost 2026). Both SAFE and RISK facilitate analysis of complex genetic interaction profiles by identifying sets of functionally related library genes that show coherent negative and/or positive genetic interactions with a selected query gene (Baryshnikova 2016; Costanzo *et al*. 2016; Billmann *et al*. 2025b; Horecka and Rost 2026). These enriched region(s) are then visualized directly on the HAP1 genetic interaction profile similarity network, enabling functional characterization of a query gene genetic interaction profile (Baryshnikova 2016; Costanzo *et al*. 2016; Horecka and Rost 2026). For example, RISK analysis using a *FANCG* query gene interaction profile as an input, showed that the “DNA replication and repair” region of the HAP1 profile similarity network is significantly enriched (*P* < 0.001) for library genes showing negative and positive interactions with *FANCG*, consistent with the general tendency of negative genetic interactions to connect functionally related genes and positive interactions to identify genes encoding members of the same nonessential pathway or protein complex (Fig. 5A)(Billmann *et al*. 2025b). The corresponding SAFE enrichment analysis can be performed by clicking on the appropriate button in the upper right corner (Fig. 5A).

**Figure 5.**
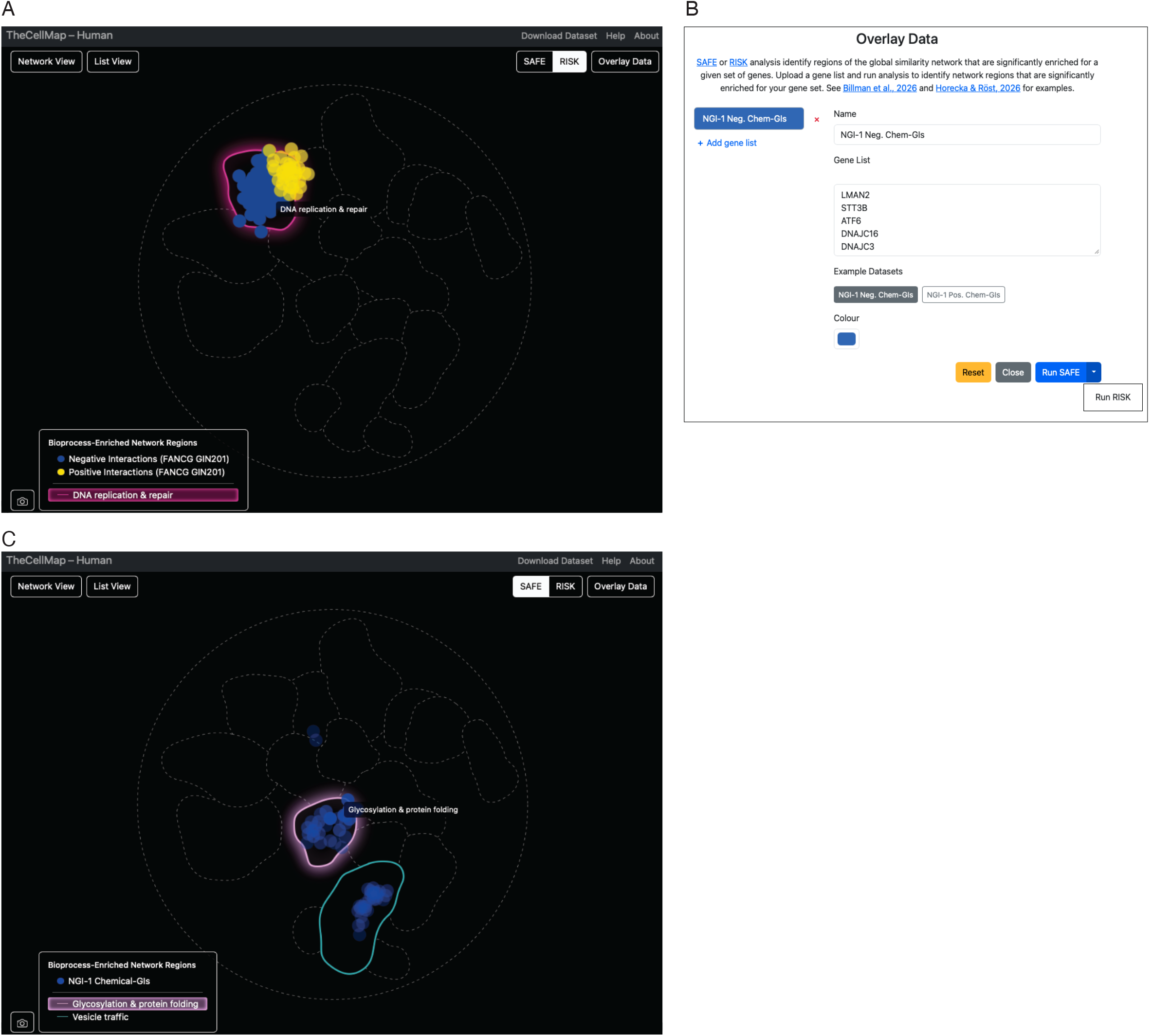
Automated SAFE analysis to functionally annotate custom gene sets. (**A**) Automated RISK or SAFE analysis (RISK shown) generates an enrichment map highlighting regions of the global HAP1 genetic interaction similarity network significantly enriched (*P* < 0.001) for library genes exhibiting negative (blue) or positive (yellow) genetic interactions with *FANCG*. The legend denotes significantly enriched network regions; functional annotations are revealed by hovering over the corresponding network regions. Users can select either RISK or SAFE analysis by clicking the appropriate button in the upper right corner. (**B**) Using the “Overlay Data” option (Figs. 1C, 4A), users may submit one or more custom gene sets for SAFE or RISK analysis, assigning each dataset a unique color. (**C**) Regions of the HAP1 profile similarity network significantly enriched (*P* < 0.001) for library genes exhibiting negative (blue) chemical-genetic interactions with NGI-1 (SAFE analysis shown). The legend indicates significantly enriched network regions, and functional annotations are revealed by hovering over the corresponding network regions. Users can run either RISK or SAFE analysis by clicking the appropriate button in the upper right corner.

### Using the HAP1 genetic interaction profile similarity network to functionally annotate gene sets

SAFE and RISK are not limited to analysis of query gene genetic interaction profiles. Both methods have been previously used in combination with yeast and human genetic interaction profile similarity networks to functionally annotate diverse gene sets derived from large-scale phenotypic screens (Baryshnikova 2016; Costanzo *et al*. 2016; Costanzo *et al*. 2021; Billmann *et al*. 2025b; Horecka and Rost 2026). From the TheCellMap.org homepage (Fig. 1C) or the List View page (Fig. 4A), clicking the “Overlay Data” button opens an interface where users can upload one or more gene sets for SAFE or RISK analysis (Fig. 5B). As an example, we uploaded a list of genes that showed negative chemical-genetic interactions in a genome-wide screen with NGI-1, a small molecule inhibitor of the oligosaccharyltransferase (OST) complex (Billmann *et al*. 2025b). Chemical-genetic interaction analysis involves profiling a small molecule against a library of mutant strains or cell lines, each with a LOF mutation in a specific gene (Parsons *et al*. 2006; Piotrowski *et al*. 2017). Mutants showing specific sensitivity to the compound define negative chemical-genetic interactions, and the collection of such genetic sensitivities constitute a chemical-genetic interaction profile that often reflects the compound’s mode-of-action (Parsons *et al*. 2006; Piotrowski *et al*. 2017). SAFE or RISK generate an outline of the HAP1 profile similarity network in a new browser tab, highlighting network regions enriched for input genes by coloring their corresponding nodes (Fig. 5C). For the NGI-1 gene set, enriched regions overlapped HAP1 network clusters corresponding to “Glycosylation & Protein Folding” and “Vesicle Traffic”, two biological processes known to be perturbed by NGI-1 exposure (Fig. 5C)(Cheng *et al*. 2022; Billmann *et al*. 2025b). Clusters significantly enriched for the input gene set are listed in the accompanying legend, and users can identify cluster identities by hovering over individual network regions or term names within the legend (Fig. 5C). SAFE and RISK enrichment maps can also be downloaded as .png files.

## Conclusion

TheCellMap.org was originally developed as an integrated platform for accessing, visualizing, and analyzing the global yeast genetic interaction network. We have expanded the resource to include systematic analysis of ~4 million human gene pairs in the human haploid HAP1 cell line, encompassing ~89,000 quantitative negative and positive genetic interactions. Mapping of human genetic interactions is ongoing, and newly generated datasets will be added to the database as they become available. In parallel, we continue to develop additional functionality, including tools for data integration and comparative analyses aimed at exploring the conservation of genetic networks from yeast to human cells. Together, these resources provide a powerful framework for uncovering phenotypic and functional relationships among genes and offer a valuable platform for gene function discovery and the study of global principles underlying genetic interaction networks.

## Acknowledgments

We thank members of the Andrews, Boone and Myers labs for comments and suggestions. This work was supported by the National Institutes of Health grants R01HG00583 (BA, CB, CLM), R01HG005084 (CLM), the Canadian Institutes of Health Research grants PJT-180285 (CB), Ontario Research Fund grants RE09-011 and RE011-006 (BA, CB), Canadian Foundation for Innovation grant 39977 (BA, CB), McLaughlin Centre Accelerator grants MC-2022-02 and MC-2024-08-02 (BA, CB), National Science Foundation grant MCB1818293 (CLM). BA holds a Tier 1 Canada Research Chair in Systematic Genetics & Cell Biology. CB is a Banting & Best Distinguished Scholar and a CIFAR Fellow in the Fungal Kingdom: Threats & Opportunities program.

